# Transient and layer-specific reduction in neocortical PV inhibition during sensory association learning

**DOI:** 10.1101/2020.04.24.059865

**Authors:** Dika A. Kuljis, Eunsol Park, Stephanie E. Myal, Claudia Clopath, Alison L. Barth

**Author notes:** **Author contributions:** Authors contributed to acquisition, analysis, experimental design, and interpretation of electrophysiological (DAK, EP, SEM, ALB), anatomical (DAK, EP, and ALB), and modeling (CC, ALB) data sets. All authors contributed to the writing of the manuscript.

## Abstract

Sensory and motor learning reorganizes neocortical circuitry, particularly manifested in the strength of excitatory synapses. Prior studies suggest reduced inhibition can facilitate glutamatergic synapse plasticity during learning, but the role of specific inhibitory neurons in this process has not been well-documented. Here we investigate whether inhibition from parvalbumin (PV)-expressing neurons is altered in primary somatosensory cortex in mice trained in a whisker-based reward-association task. Anatomical and electrophysiological analyses show PV input to L2/3, but not L5, pyramidal (Pyr) neurons is rapidly suppressed during early stages of sensory training, effects that are reversed after longer training periods. Importantly, sensory stimulation without reward does not alter PV-mediated inhibition. Computational modeling indicates that reduced PV inhibition in L2/3 selectively enables an increase in translaminar recurrent activity, also observed during SAT. PV disinhibition in superficial layers of the neocortex may be one of the earliest changes in learning-dependent rewiring of the cortical column.

**Impact statement:** Tactile learning is associated with reduced PV inhibition in superficial layers of somatosensory cortex. Modeling studies suggest that PV disinhibition can support prolonged recurrent activity initiated by thalamic input.

## Introduction

Disinhibition of cortical circuits during learning is associated with increased pyramidal (Pyr) neuron activity, excitatory synaptic plasticity, and the formation of memory-specific ensembles (Letzkus et al., 2015). Evidence for decreased inhibition has been observed acutely during task engagement, and also as structural and functional changes that persist beyond task engagement during the early stages of learning. In humans, acute reductions in GABA signaling during motor task acquisition are positively correlated with motor learning (Floyer-Lea et al., 2006; Stagg et al., 2011), while in mice, neocortical layer 2/3 (L2/3) parvalbumin (PV) cell activity is acutely suppressed in response to stimulus presentation during auditory fear conditioning (Letzkus et al., 2011). Additionally, more persistent functional and structural changes to inhibition have also been observed in rodents. During early stages of learning, the frequency of inhibitory postsynaptic currents (IPSCs) onto L2/3 Pyr cells is reduced during early stages of auditory and motor learning (Sarro et al., 2015; Kida et al., 2016), and alterations to PV and somatostatin (SST) axonal boutons during motor learning have also been observed in superficial layers (Donato et al., 2013; Chen et al., 2015b). It has been hypothesized that disinhibition facilitates the rewiring of cortical networks during learning (Letzkus et al., 2015; Williams and Holtmaat, 2019), but key mechanistic details such as the specific inhibitory cell types and targets involved, the stage during the learning trajectory when disinhibition occurs, and the persistence of disinhibition all remain unclear.

PV-expressing fast-spiking interneurons are the most abundant type of interneuron in cortex and the most potent source of inhibition onto Pyr neurons (Markram et al., 2004; Pfeffer et al., 2013). They play a critical role in sensory-evoked feedforward and feedback inhibition, as well as sensory processing (Packer and Yuste, 2011; Jiang et al., 2015; Barth et al., 2016; Audette et al., 2017; Li et al., 2019). These functions are mediated by the broad distribution of their inhibitory synapses on Pyr axons, soma, and both proximal and distal dendrites (Kubota et al., 2015; Tremblay et al., 2016; Kuljis et al., 2019). Transient reductions in PV inhibition of cortical Pyr neurons have been characterized during passive manipulation of sensory input (Hengen et al., 2013; Kuhlman et al., 2013; Kaplan et al., 2016; Gainey et al., 2018; Cisneros-Franco and de Villers-Sidani, 2019), but there is also a growing appreciation for the role of PV plasticity in sensory and motor learning (Donato et al., 2013; Chen et al., 2015b; Letzkus et al., 2015). Critical gaps in our knowledge of PV plasticity during learning include whether it is manifested as structural synaptic plasticity or a change in PV intrinsic excitability, its target selectivity, and whether it can differentiate between passive sensory experience and reward-based learning.

Here we use an automated, homecage system for sensory association training (SAT) to examine the laminar location and trajectory of changes in PV-mediated inhibition in primary somatosensory (barrel) cortex during learning. Our prior studies have shown that excitatory synaptic strengthening sequentially progresses across the cortical column during SAT, starting in deep layers and progressing to superficial layers (Audette et al., 2019). In accordance with the disinhibition hypothesis (Letzkus et al., 2015), we predicted that PV disinhibition would be initiated in deep layers, where thalamic input potentiation is first observed, and would then proceed to superficial layers prior to the emergence of excitatory strengthening there. Instead, we observed a rapid suppression of PV inhibition of Pyr neurons in superficial layers and no alteration in deep layers at any time point. This layer-specific reduction in inhibitory input is transient, since PV inhibition returns to control levels after 5 days of SAT. Anatomical analyses show that the reduction in PV input was specifically linked to the post-synaptic removal of PV-synapses on L2/3 Pyr neurons. Computational modeling indicates that a reduction of L2/3 PV inhibition can facilitate stimulus-evoked recurrent activity across layers. Importantly, PV inhibitory plasticity was not generated following sensory stimulation alone, indicating that PV neurons are part of a neural circuit that can differentiate reward-association training from passive sensory input.

## Results

### Prolonged sensory association training reveals multiple stages of learning

We used an automated, home-cage training environment to train freely-moving mice for whisker-stimulus association learning as previously described (Audette et al., 2019; Figure 1). Our prior studies indicate that SAT drives progressive changes in anticipatory licking and excitatory synaptic strength that are initiated early in the training period (Audette et al., 2019). Initially we wanted to determine the timecourse of changes in anticipatory licking as a measure of association learning across a prolonged training period, in order to select discrete timepoints for analysis of cortical disinhibition. To quantify behavioral changes, the rate of anticipatory licking (300 ms prior to water delivery) for water and blank trials was compared across the training period. Control mice were housed in the same chamber, but without whisker stimulation coupled to water delivery.

**Figure 1.**
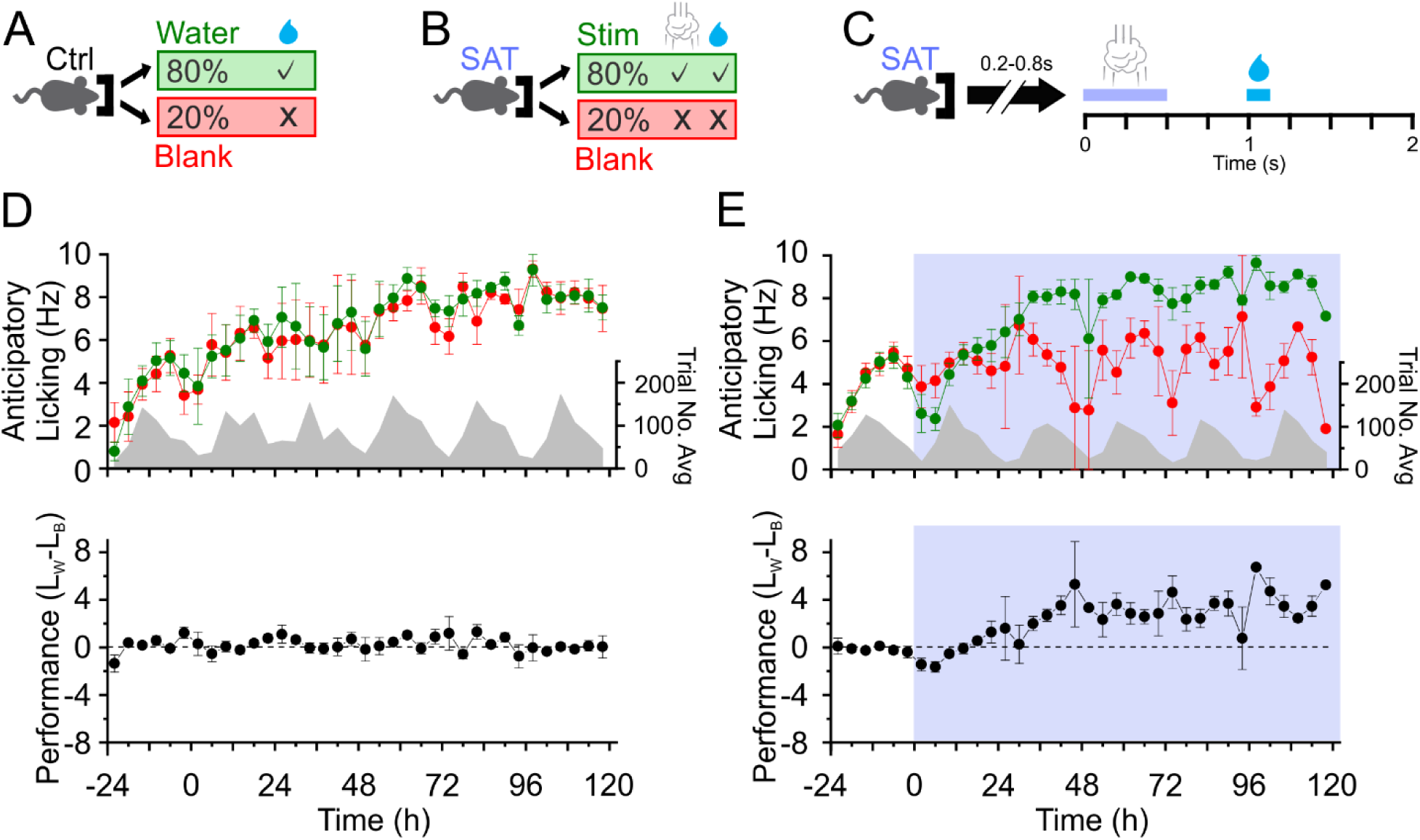
Prolonged sensory association training (SAT) reveals multiple stages of learning. **(A)** During the acclimation period, control animals receive water on 80% of trials. **(B)** On the onset of SAT, animals receive a gentle airpuff whisker stimulus (500 ms; 6psi) prior to water delivery 1 sec after airpuff onset on stimulation (stim) trials. (**C**) Schematic of trial structure. Nose-poke triggers random delay prior to trial onset. (**D)** Top: mean anticipatory lick rate for Ctrl water (green) and blank (red) trials. Grey, the distribution of average trial number over time. Bottom: mean performance (L_w_-L_b_; see methods)). Ctrl24, n=14 animals; Ctrl120, n=5 animals. **(E)** As in ***D***, but for SAT mice. SAT trials shaded in blue. SAT24, n=19 animals; SAT120, n=5 animals.

Control animals did not show differences between lick frequencies for water-delivery versus “blank” trials, as there was no predictive stimulus that would enable animals to differentiate these two trials (Figure 1A-D). Similar to our previous results (Audette et al., 2019), SAT animals exhibited a progressive increase in stimulus-cued anticipatory licking behavior across the first training day (Figure 1E). At the end of the first day of training, lick rates to water versus blank trials were greater on average (Lick_Water_ 5.8±2.5 versus Lick_Blank_ 5.1±2.4 Hz), but there was substantial heterogeneity in animal performance, where slightly more than half of the animals (11/19) showed increased anticipatory licking on water versus blank trials. By the second day of training, anticipatory lick rates on water trials consistently exceeded those of blank trials, an indication that animals had learned the association (see also Audette et al., 2019). After five days of training, changes in anticipatory lick rates to both stimulus and blank trials had plateaued and all trained animals (5/5) showed a significant difference in stimulus-associated anticipatory licking (Lick_Water_ 8.5±0.9 versus Lick_Blank_ 5.7±1.5 Hz; Figure 1E). Based on these results, we selected the 24 hour (SAT24) and five day (SAT120) timepoints to reflect early and late learning to investigate the role of PV disinhibition in SAT-related plasticity within the barrel cortex.

### Sensory association training induces a transient, layer-specific reduction of PV inhibition of Pyr neurons

To determine whether direct PV input to neocortical neurons was altered during SAT, we used brain tissue from PV-Cre × Ai32 transgenic mice for optogenetic analysis of PV-IPSC amplitude in Pyr neurons. L2/3 and L5 Pyr neurons were targeted in series for whole-cell patch-clamp recordings in acute brain slices from control and trained mice (Figure 2). Light-evoked PV-mediated IPSCs (PV-IPSCs) were recorded by holding the post-synaptic Pyr cell at −50 mV. After just 24 hrs of training, mean PV-IPSC amplitude was suppressed by ~35% in L2/3 Pyr neurons, a difference that was highly significant (24 hrs SAT 251±34 versus control 377±28 pA, *p*=0.003; Figure 2A-C, Figure 2-Source data 1). Importantly, light-evoked currents were abolished by the application of the GABA_A_-receptor antagonist picrotoxin, indicating that they were solely generated by inhibitory PV neurons (Figure 2C_1_).

**Figure 2.**
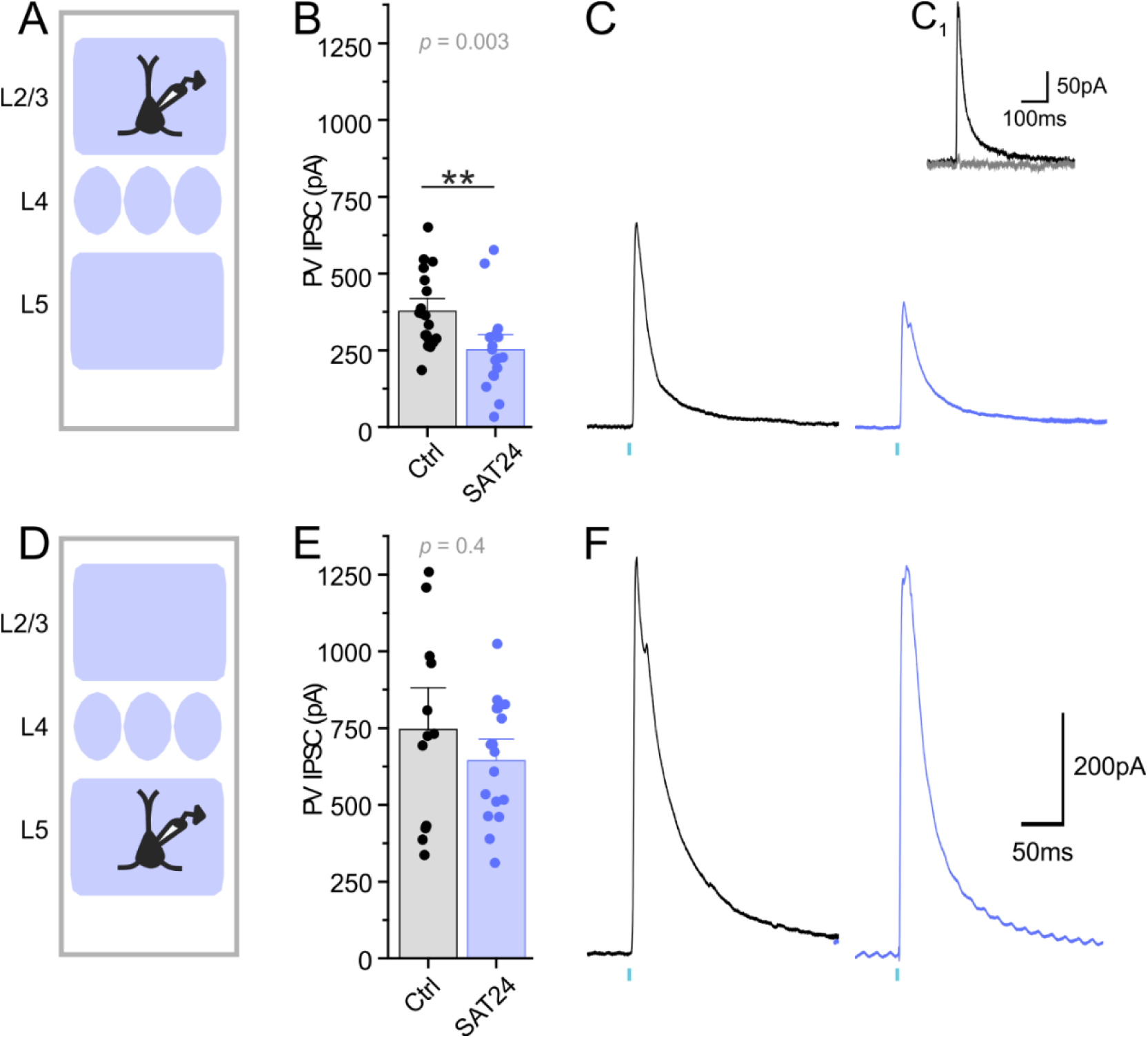
Reduced PV inhibition in supragranular Pyr neurons following 24 hrs SAT. (**A**) Schematic of L2/3 Pyr neuron targeting in PV-Cre × Ai32 mice. (**B**) PV-IPSC amplitude for L2/3 neurons from Ctrl (black) and SAT24 (blue) animals. L2/3: Ctrl n=19 cells, 3 animals; SAT24 n=17 cells, 5 animals. (**C**) Representative PV-IPSC from Ctrl (black) and SAT24 (blue) L2/3 Pyr neuron following stimulation (blue tick mark). (**C_1_**) PV-IPSC recorded before (black) and after bath application of picrotoxin (grey). (**D**) Schematic of L5 Pyr targeting. (**E-F**) As in ***B*** and ***C***, but for L5 Pyr neurons. L5: Ctrl n=13 cells, 4 animals; SAT24 n=17 cells, 5 animals.

Reduced PV-IPSC amplitude in L2/3 Pyr neurons could come from changes in postsynaptic receptor properties or decreased presynaptic release probability. To assess this, we compared the paired-pulse ratio (PPR; amplitude of peak 2/peak 1) of PV-IPSCs in response to paired light pulses (150 ms interstimulus interval). For L2/3 Pyr neurons, PPR appeared unchanged after SAT compared to controls (24 hrs SAT 0.58±0.04 versus control 0.59±0.04, *p*=0.9, n=4), suggesting presynaptic plasticity does not underlie reduced PV inhibition in L2/3 Pyr neurons.

Unlike L2/3, L5 Pyr neurons did not show a significant SAT-dependent reduction in PV-IPSC amplitude (24 hrs SAT 644±47 versus control 745±91 pA, *p*=0.4; Figure 2D-F, Figure 2-Source data 1). The small decrease in mean PV-IPSC amplitude in L5 Pyr neurons at 24 hrs SAT raised the possibility that PV disinhibition might have been rapidly induced at the onset of SAT, but had begun to renormalize by the 24-hour timepoint. To test this, mice underwent SAT for 12 hrs and PV-IPSC amplitude in L5 Pyr neurons was assessed. However, at this earlier timepoint, mean PV-IPSC amplitude was virtually identical to control levels (12 hrs SAT 768±89 versus control 745±91 pA, *p*=0.7; SAT12 n=12 cells, 3 animals; data not shown). In contrast, PV-IPSCs in L2/3 Pyr neurons at 12 hrs of SAT already appeared somewhat reduced (12 hrs SAT 283±65 versus control 377±28 pA, *p*=0.1; SAT12 n=8 cells, 3 animals; data not shown). Overall, these findings suggest PV input suppression is rapid, pronounced, and concentrated on L2/3 Pyr neurons.

Is PV disinhibition stable across the learning trajectory? Our prior work indicates that excitatory synaptic changes, particularly in L2/3 Pyr neurons, progressively increase with longer training periods (Audette et al., 2019). However, after 5 days of SAT, mean PV-IPSC amplitude in L2/3 Pyr neurons reverted to baseline values and were similar to age-matched controls (120 hrs SAT 365±22 versus control 355±37 pA, *p*=0.5; Figure 3A-C, Figure 3-Source data 1). Mean PV-IPSC amplitude in L5 Pyr neurons was again unchanged compared to baseline values (120 hrs SAT 619±59 pA versus control 609±50 pA, *p*=1.0; Figure 3D-F, Figure 3-Source data 1). L5 Pyr neurons are comprised of a heterogeneous class of Pyr neurons defined by morphology, firing phenotype, and axonal target (Lee et al., 2014; Kim et al., 2015). Analysis of PV input to regular-spiking and intrinsically bursting L5 Pyr neurons across SAT timepoints did not suggest selective regulation of PV input (data not shown). Overall, these findings suggest SAT rapidly initiates a reduction in PV input to Pyr neurons, specifically targeted to L2/3 but not L5 Pyr neurons, and that these changes are restricted to the early stages of SAT.

**Figure 3.**
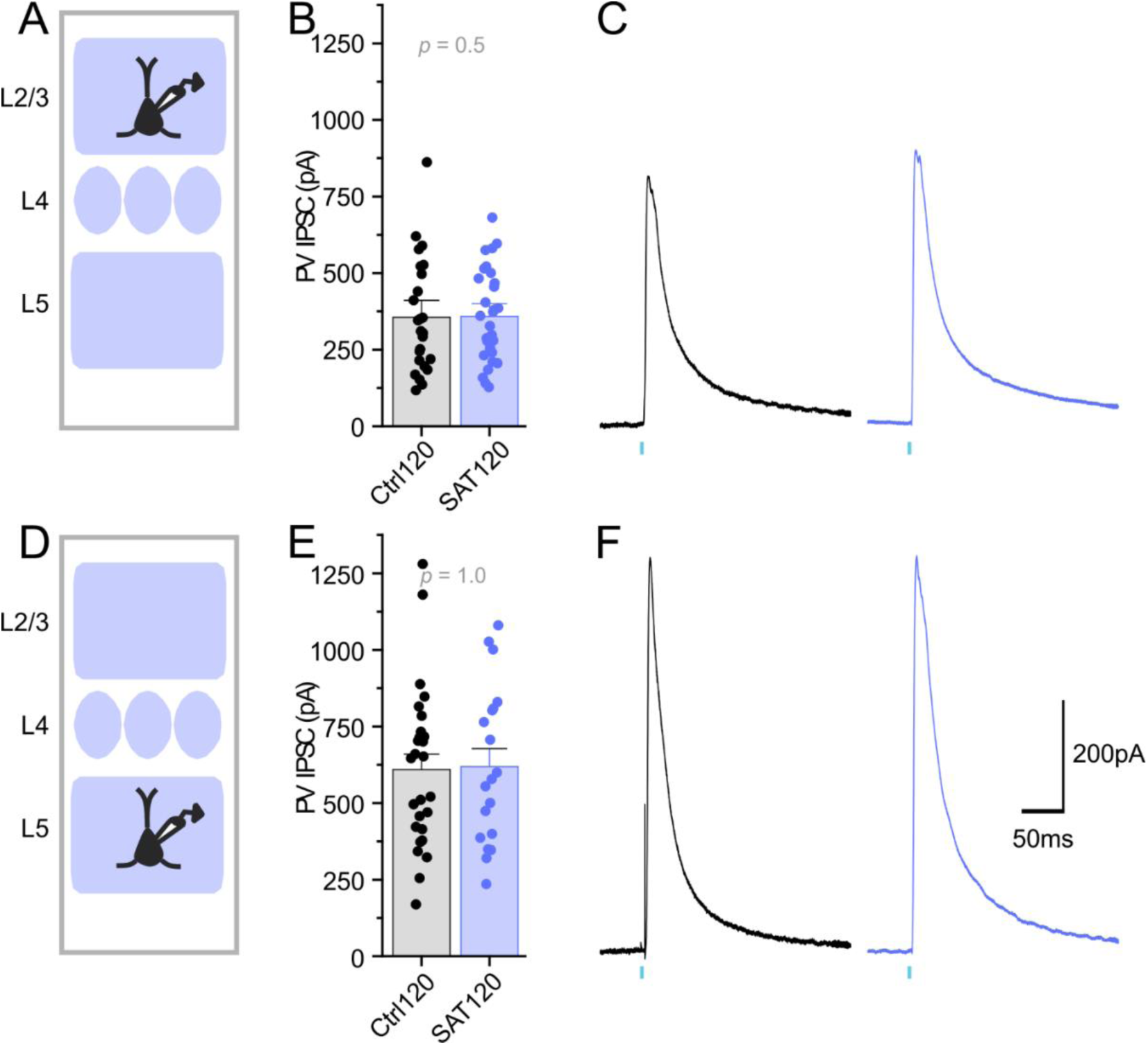
PV inhibition of supragranular Pyr neurons is restored following 120 hrs of SAT. (**A**) Schematic of L2/3 Pyr neuron targeting in PV-Cre × Ai32 mice. (**B**) PV-IPSC amplitude for L2/3 neurons from Ctrl120 (black) and SAT120 (blue) animals. L2/3: Ctrl120 n=25 cells, 5 animals; SAT120 n=40 cells, 4 animals. (**C**) Representative Ctrl120 (black) and SAT120 (blue) PV-IPSC recorded in L2/3 Pyr neuron following stimulation (blue tick mark). (**D**) Schematic of L5 Pyr targeting. (**E-F**) As in ***B-C***, but for L5 Pyr neurons. L5: Ctrl120 n=27 cells, 5 animals; SAT120 n=19 cells, 4 animals.

### Passive sensory experience alone does not alter PV inhibition of Pyr neurons

Our prior studies showed that passive sensory stimulation in the absence of reward was not sufficient to potentiate thalamocortical inputs to neocortical neurons (Audette et al., 2019). To determine whether PV disinhibition was unique to reward-association training or could be induced by exposure to the sensory stimulus alone, we adjusted the trial structure so that water delivery was uncorrelated with the multiwhisker stimulus, a paradigm referred to as pseudotraining (Figure 4A). As expected, pseudotrained animals showed no difference in anticipatory licking between stimulation and blank trials (Figure 4B). Mean PV-IPSC amplitude in L2/3 neurons from pseudotrained mice was not significantly different from control (24 hrs pseudotraining 427±29 versus control 356±30 pA, *p*=0.1; Figure 4C-E, Figure 4-Source data 1). L5 Pyr neurons also showed no change in PV-IPSCs (24 hrs pseudotraining 573±84 versus control 651±65 pA, *p*=0.5; Figure 4F-H, Figure 4-Source data 1). Thus, sensory stimulation in the absence of reward is not sufficient to drive a reduction in PV inhibition in either L2/3 or L5.

**Figure 4.**
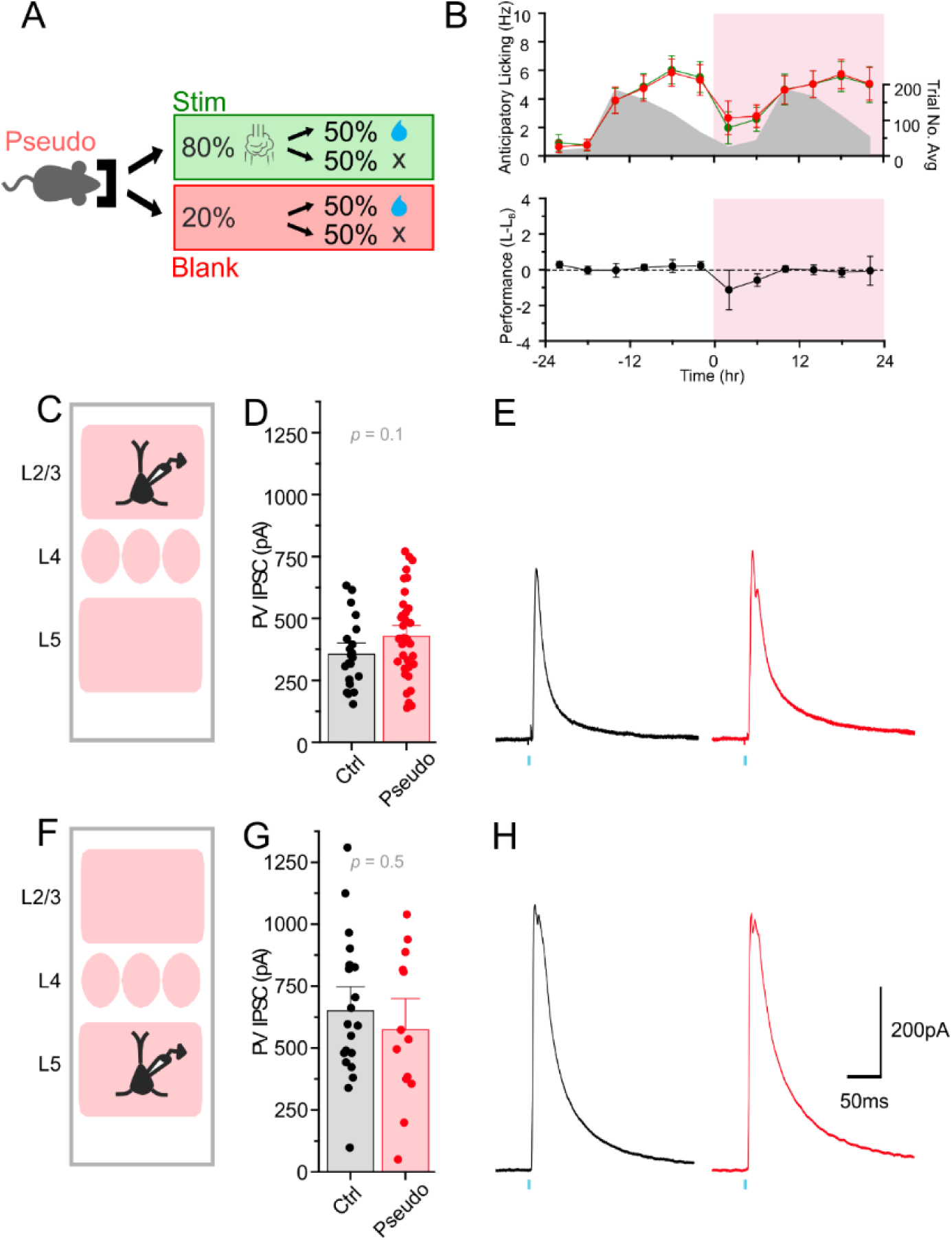
Reward-uncoupled pseudotraining does not affect PV inhibition of Pyr neurons. (**A**) Schematic of training conditions. Pseudotrained animals receive airpuff stimulation (stim) for 80% of trials, and no stim on 20% of blank trials. 50% of both stim and blank trials receive water reward. (**B**) Top: anticipatory lick rate for pseudotrained animals on stim (green) and blank (red) trials. Grey, the distribution of average trial number over time. Bottom: average performance (L_w_-L_b_) of pseudotrained animals. Pseudotraining onset shaded in pink. (**C**) Schematic of L2/3 Pyr neuron targeting in PV-Cre × Ai32 mice. (**D**) PV-IPSC amplitude for L2/3 neurons from Ctrl (black) and Pseudo (red) animals. L2/3: Ctrl (pseudo) n=21 cells, 4 animals; Pseudo24 n=31 cells, 4 animals. (**E**) Representative PV-IPSC from Ctrl (black) and Pseudo (red) L2/3 Pyr neuron following stimulation (blue tick mark). (**F**) Schematic of L5 Pyr targeting. (**G-H**) As in ***D*** and ***E***, but for L5 Pyr neurons. L5: Ctrl (pseudo) n=20 cells, 4 animals, Pseudo n=13 cells, 3 animals.

### Sensory association training effects on intrinsic membrane properties of Pyr and PV neurons

Neural circuit plasticity and homeostasis is a complex process that can involve alterations to postsynaptic neuron excitability in addition to synaptic strength, and decreased Pyr neuron excitability could offset the network consequences of decreased PV synaptic drive through homeostatic mechanisms. Similar to our previously published findings (Audette et al., 2019), we did not find SAT-dependent differences in current-evoked firing for either L2/3 or L5 Pyr neurons (data not shown). These findings suggest alterations in Pyr neuron excitability would not offset reduced PV synaptic drive.

Alternatively, an increase in PV neuron excitability could compensate for apparent reductions in the ChR2-evoked IPSC that could offset synaptic effects during network activity. We thus examined the intrinsic excitability and electrophysiological properties of PV neurons using whole-cell current-clamp recordings. Neither resting membrane potential, input resistance, the number of optically-evoked spikes, rheobase current, nor current-evoked firing were different in PV neurons after 24 hrs of training (Figure 5, Figure 5-Source data 1). Overall, these findings suggest that reduced PV-mediated inhibition of L2/3 Pyr neurons early during SAT is likely to be manifest during network activation, and cannot simply be explained by a decrease in PV neuron excitability.

**Figure 5.**
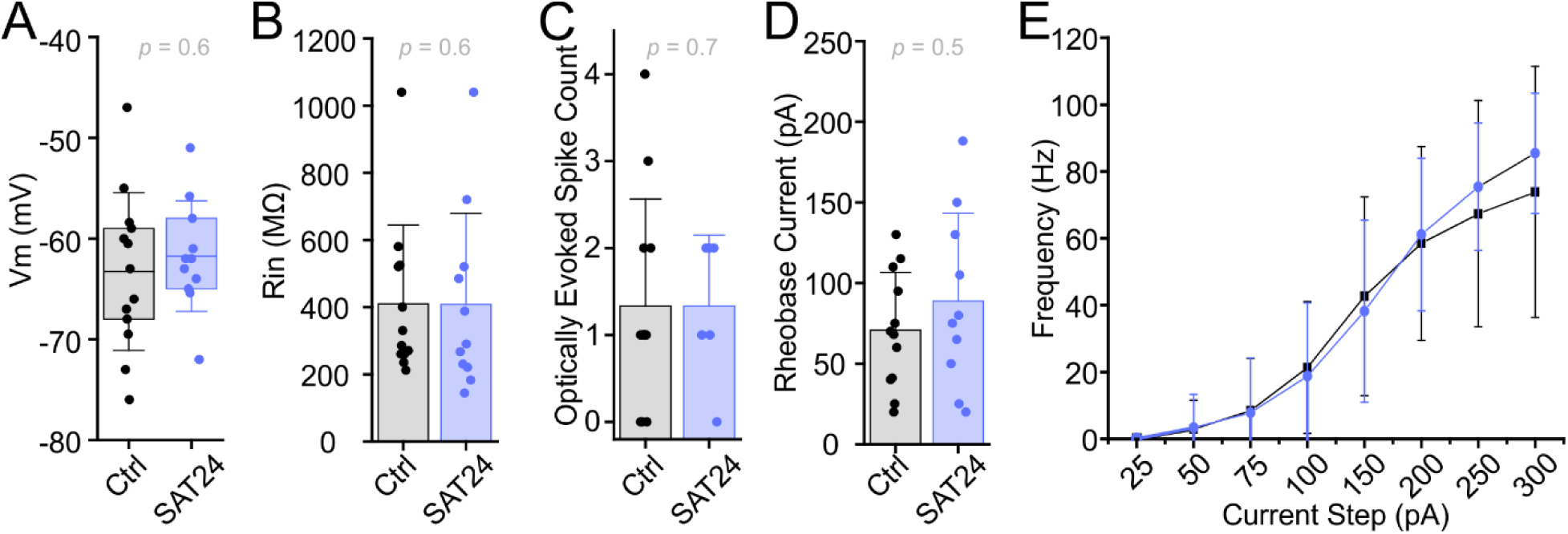
L2/3 PV neuron excitability is unchanged after 24 hrs of SAT. (**A**) Membrane potential (*V*_*m*_) of Ctrl (black) and SAT24 (blue) PV neurons. Box is 25^th^ and 75^th^ quartile, whiskers are SD, and midline is mean. Ctrl, n=13 cells, 4 animals; SAT24, n=11 cells, 6 animals. (**B**) Input resistance (R_in_). Ctrl n=12 cells, 3 animals; SAT24 n=11 cells, 6 animals. (**C**) Optically evoked spike count. Ctrl n=12 cells, 3 animals; SAT24 n=6 cells, 2 animals. (**D**) Rheobase current. Ctrl n=12 cells, 3 animals; SAT24 n=10 cells, 5 animals. (**E**) No effect of 24 hrs of SAT on PV neuron firing rate responses to positive current injection steps (ANOVA_SAT_: F_(1,186)_=0.82, *p*=0.37; ANOVA_SATxStep_: F_(7,186)_=0.38, *p*=0.92). Ctrl n=9 cells, 4 animals; SAT24 n=9 cells, 6 animals. Line and dot plot represents mean±SD.

### Sensory association training induced structural plasticity

To determine whether transiently-reduced PV inhibition of L2/3 Pyr neurons during SAT was associated with pre- and/or postsynaptic anatomical plasticity of PV synapses, we deployed fluorescence-based quantitative synapse analysis using a neuroligin-based synaptic tagging molecule (FAPpost), previously shown to detect PV synapses with high accuracy (Kuljis et al., 2019). Postsynaptic Pyr neurons were virally transduced with a cell-filling dTomato (dTom) and postsynaptic FAPpost in PV-Cre × Ai3 transgenic mice for comprehensive YFP labeling of PV neurites. Confocal imaging and digital alignment of presynaptic PV structures with postsynaptic sites on target Pyr neurons was used to examine the distribution of PV-assigned FAPpost puncta (PV synapses) on soma and dendrites for a target Pyr neuron.

Use of postsynaptic molecular markers in conjunction with presynaptic neurite localization can provide an accurate way to detect and quantitate the number of input-specific synapses (Kubota et al., 2015; Kuljis et al., 2019; Figure 6 Supplement 1). For L2/3 Pyr neurons after 24 hrs of SAT, PV synapse density was lower on both dendrites and soma (dendrites 24 hrs SAT 0.18±0.14 versus control 0.29±0.15 per µm; soma 24 hrs SAT 0.38±0.39 versus control 0.65±0.30 per 10µm^2^; Figure 6A-G, Figure 6-Source data 1). This reduction was similar in magnitude to the decrease observed through electrophysiological measurements, approximately 35%.

**Figure 6.**
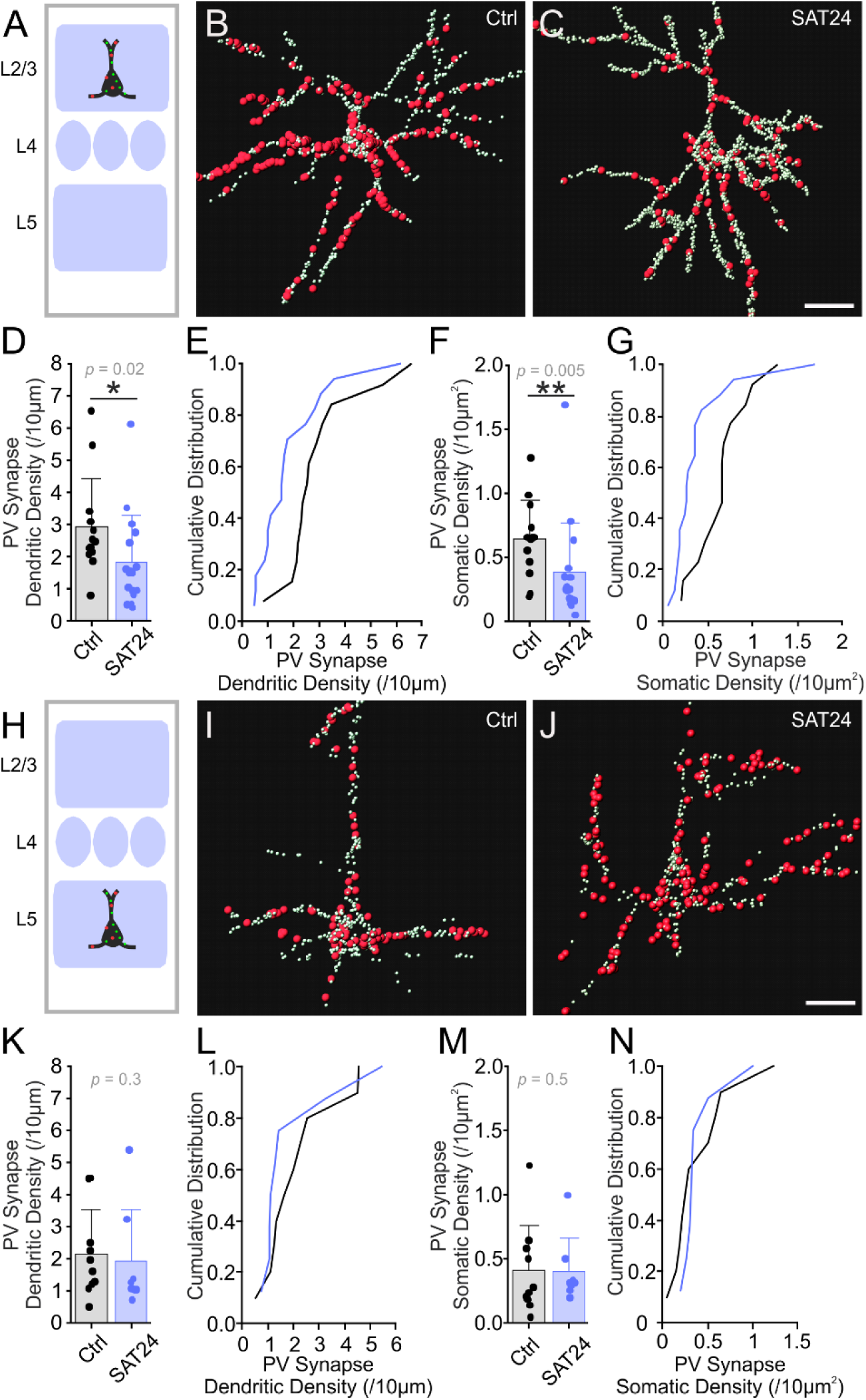
SAT reduces PV synapse density in L2/3 but not L5 Pyr neurons. (**A**) Schematic of L2/3 Pyr anatomical analysis. (**B**) Representative Ctrl L2/3 Pyr neuron with PV-assigned (large red) and unassigned (small green) FAPpost-labeled synapses. (**C**) As in ***B***, but for a L2/3 Pyr neuron after 24 hrs SAT. (**D**) Mean L2/3 Pyr dendritic PV synapse density. (**E**) Cumulative frequency distribution for dendritic PV synapse density for a L2/3 Pyr neuron in Ctrl (black) and SAT24 (blue). (**F-G**) As in ***D-E***, but for somatic PV synapse density. L2/3: Ctrl n=17 cells, 5 animals; SAT24 n=17 cells, 5 animals. (**H-N**) As in ***A-G***, but for L5 Pyr neurons. L5: Ctrl n=10 cells, 4 animals; SAT24 n=8 cells, 5 animals. Scale bar = 20µm.

In contrast, L5 Pyr PV synapse density was unchanged for both dendrites and soma (dendrites 24 hrs SAT 0.19±0.16 versus control 0.22±0.14 per µm; soma 24 hrs SAT 0.41±0.26 versus control 0.41±0.35 per 10µm^2^; Figure 6H-N, Figure 6-Source data 1), consistent with PV-IPSC measurements. Overall, these findings suggest that postsynaptic structural plasticity underlies reduced PV inhibition of L2/3 Pyr neurons observed early during SAT.

Postsynaptic plasticity may occur at the same time as presynaptic structural plasticity. To test whether decreased PV inhibition of Pyr neurons was accompanied with the loss of presynaptic PV+ terminals, we also compared the density of PV-neurite associations across the dendrites and soma of individual Pyr neurons (Figure 7). For L2/3 Pyr neurons, PV-neurite associations along dendrite and soma were unchanged after SAT (dendrite 24 hrs SAT 1.2±0.3 versus control 1.4±0.6 per µm; soma 24 hrs SAT 1.7±0.7 versus control 1.8±0.8 per 10µm^2^; Figure 7D-G, Figure 7-Source data 1). For L5 Pyr neurons, the density of PV-neurite associations along dendrites and soma was also similar across conditions (dendrites 24 hrs SAT 2.0±0.6 versus control 2.1±0.4 per µm; soma 24 hrs SAT 2.0±0.5 versus control 2.1±0.8 per 10µm^2^; Figure 7K-N, Figure 7-Source data 1). These data suggest that decreased PV inhibition in L2/3 Pyr neurons is not accompanied by the large-scale retraction of PV terminals.

**Figure 7.**
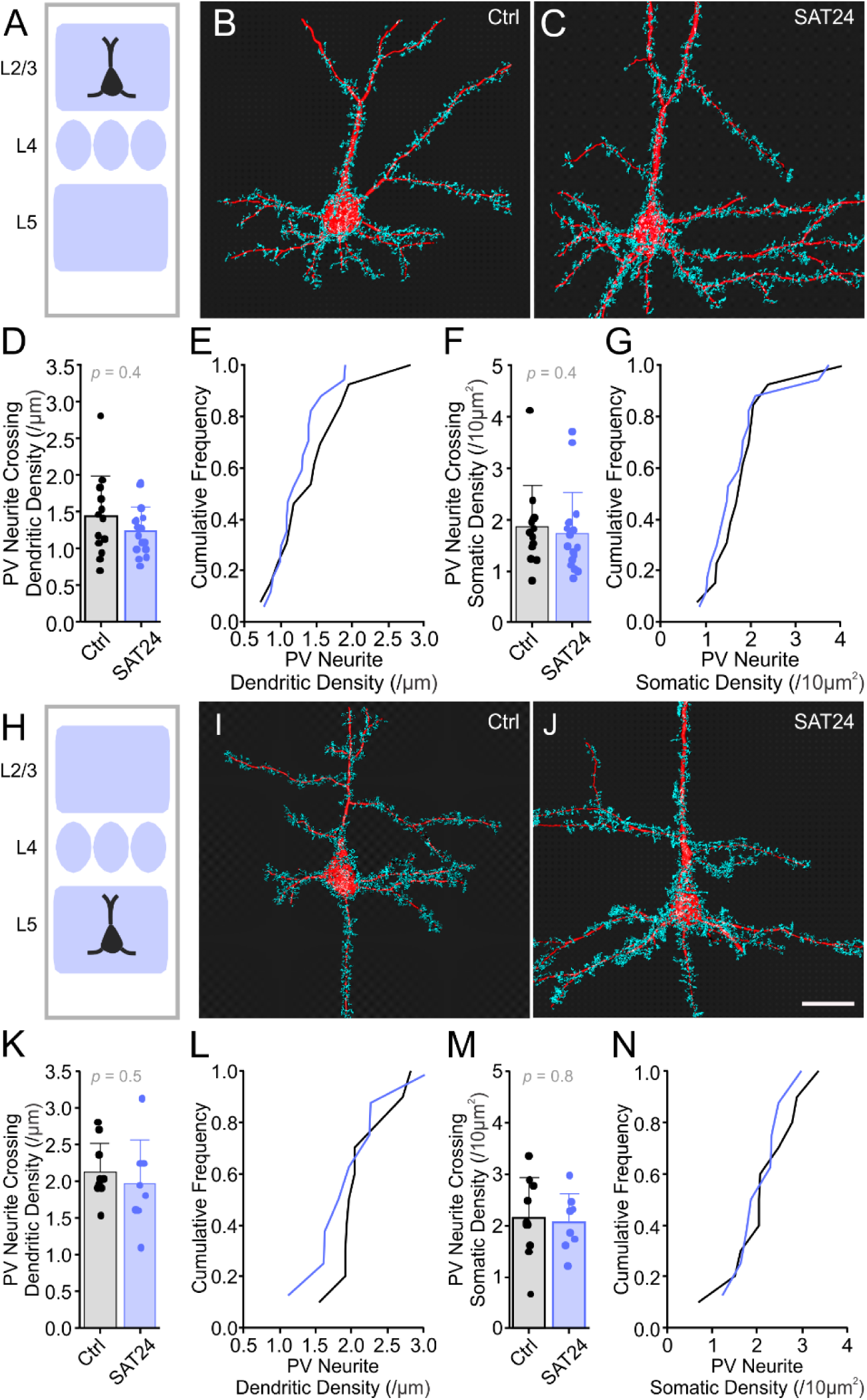
SAT does not alter presynaptic PV neurite association with L2/3 and L5 Pyr neurons. (**A**) Schematic of L2/3 Pyr anatomical analysis. (**B**) Representative Ctrl L2/3 Pyr neuron (red) and associated presynaptic PV neurites (blue). (**C**) As ***B***, but for a L2/3 Pyr neuron after SAT. (**D**) Mean density of PV neurite association on L2/3 Pyr dendrites. (**E**) Cumulative frequency distribution of PV neurite association density along dendrites of L2/3 Pyr neurons in Ctrl (black) and SAT24 (blue). (**F**) Mean density of PV neurite association on L2/3 Pyr soma. (**G**) As in ***E***, but for somatic PV neurite associations. L2/3: Ctrl n=17 cells, 5 animals; SAT24 n=17 cells, 5 animals. (**H-N**) As in ***A-G***, but for L5 Pyr neurons. L5: Ctrl n=10 cells, 4 animals, SAT24 n=8 cells, 5 animals. Scale bar = 20µm.

### L2/3 disinhibition specifically regulates recurrent cortical network activity

In somatosensory cortex, L2/3 Pyr neurons exhibit sparse firing for both spontaneous activity and also sensory-evoked responses (Barth and Poulet, 2012), a phenomenon that is at least partially due to strong feedback inhibition from PV neurons (Jouhanneau et al., 2018). To investigate how SAT-dependent reductions in feedback inhibition from PV neurons would impact thalamically-evoked network activity, we developed a computational model to isolate and compare the effects of PV input changes in L2/3 and L5, key targets of early SAT-dependent plasticity (Audette et al., 2019). We focused on activity generated by the posterior-medial nucleus of the thalamus (POm), since this pathway is selectively enhanced by SAT (Audette et al., 2019).

Experimental measurements of input strength for POm and PV synapses onto L2/3 and L5 Pyr neurons were used to construct the model (Audette et al., 2017). Importantly, experimental data indicate that L5 but not L2/3 PV neurons receive direct synaptic input from POm (Audette et al., 2017). The small circuit we constructed also included reciprocal connectivity between L2/3 and L5 Pyr neurons (Jiang et al., 2015; Lefort and Petersen, 2017), as well as an increase in POm synaptic strength onto L5 but not L2/3 Pyr neurons, as has been described in the initial stages of SAT (Audette et al., 2019).

Our prior studies in acute brain slices indicate that after just 24 hrs of SAT, both L2/3 and L5 Pyr neurons show a significant increase in firing both during thalamic (POm) stimulation and also in the post-stimulus window (Audette et al., 2019). To determine whether reduced L2/3 PV inhibition was sufficient to enable recurrent activity, we created a simple biologically-grounded model of a multi-layered cortical network with feedforward and feedback PV inhibition in L5 and feedback inhibition in L2/3 and POm drive to both layers (Figure 8A). Similar to experimental data from control animals, POm stimulation did not drive prolonged post-stimulus activity across L2/3 and L5 (Audette et al., 2019).

**Figure 8.**
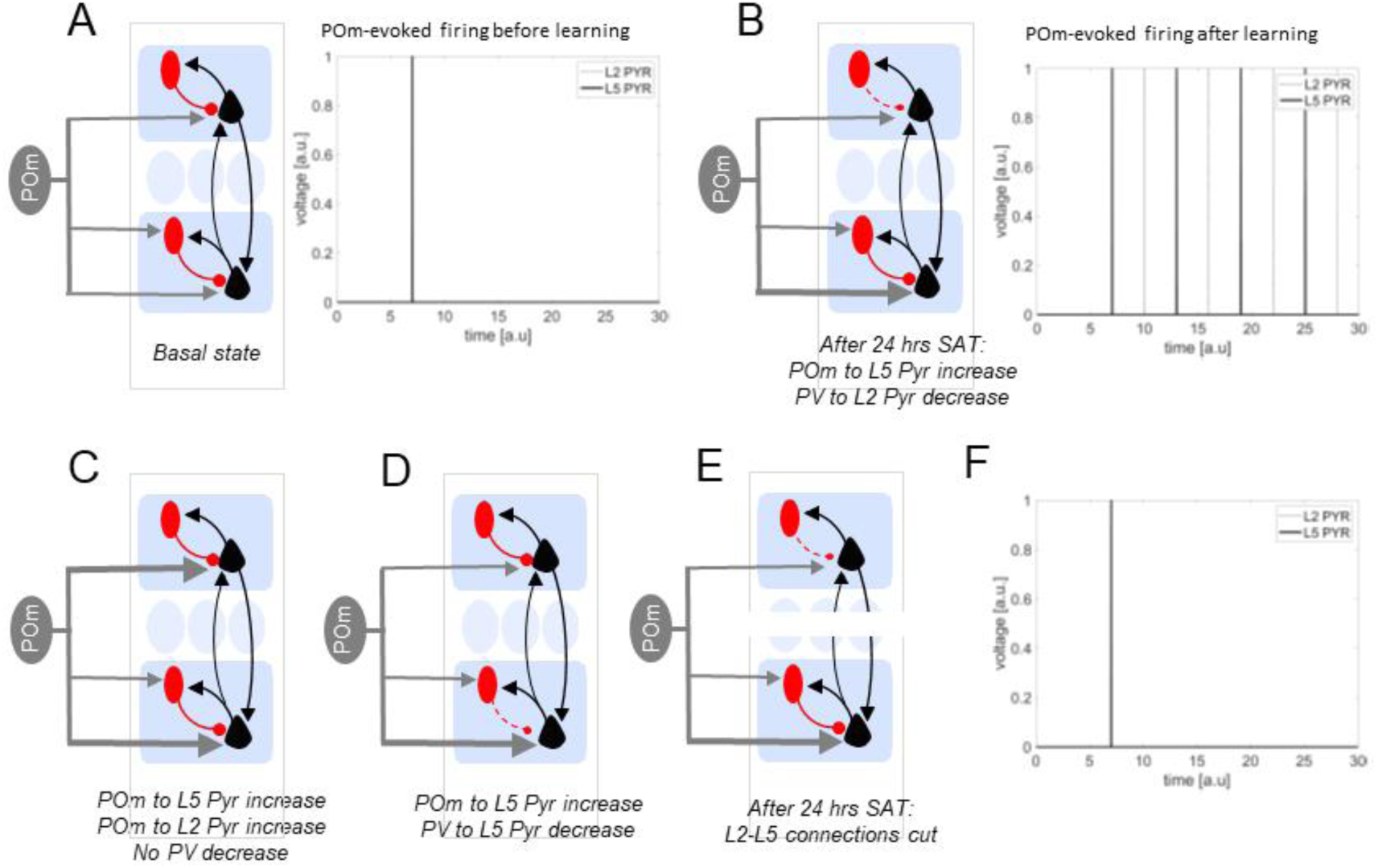
Computational model shows L2-specific PV disinhibition is sufficient to generate recurrent activity across L2/3 and L5. **(A**) Left, schematic of basal synaptic inputs included in model. POm inputs in grey, Pyr neurons in black, and PV neurons in red. Right, output from integrate-and-fire model of POm-evoked firing of L5 Pyr (black) and L2/3 Pyr (dotted) with a single POm stimulus at time=6 au. Black line at t=7 indicates firing of L5 Pyr. (**B**) Left, schematic of synaptic weights adjusted to match changes at 24 hrs SAT. Right, as in ***A*** but where POm input to L5 Pyr is strengthened by 20% and PV feedback to L2/3 Pyr is reduced by 40%. L5 Pyr firing precedes L2/3 Pyr as before, but now there is reciprocal excitation across layers that can escape feedback PV inhibition. (**C**) Schematic, model where only POm input strength is increased, but to both L2/3 and L5 Pyr. POm stimulation is not sufficient to drive recurrent L5-L2/3 activity; see ***F***. (**D**) Schematic, model where POm input to L5 is increased and feedback inhibition from PV to L5 Pyr is reduced. POm stimulation is not sufficient to drive recurrent L5-L2/3 activity; see ***F***. (**E**) Schematic, model as in ***B**,* but where L2/3-L5 Pyr connections are removed. POm stimulation does not drive recurrent L5-L2/3 activity; see ***F***.

After 24 hrs of SAT, POm inputs to L5 Pyr are ~20% larger (Audette et al., 2019) and PV input in L2/3 is reduced by ~35% (Figure 2, 6). Adjusting these values in the model circuit revealed that brief POm stimulation now initiated a prolonged recurrent excitatory loop between L2/3 and L5 (Figure 8B). Increasing POm input strength to L5 Pyr or to both L2/3 and L5 Pyr neurons, as occurs after longer periods of SAT (Audette et al., 2019) without reducing L2/3 PV feedback inhibition was similar to control conditions, with no sustained excitation across layers (Figure 8C, F). Thus, reduced PV inhibition to L2/3 neurons is critical for the generation of POm-evoked recurrent activity within the cortical circuit.

Decreasing PV input to L5 Pyr neurons had no effect on recurrent excitation, in part due to the strong feedforward POm drive onto L5 PV neurons (Figure 8D, F). Indeed, even when feedback inhibition from L5 PV neurons was eliminated, POm activation was still not able to initiate recurrent activity between L2/3 and L5, underscoring the role of feedback PV inhibition in L2/3. Finally, modeling analysis showed that prolonged firing required interaction between L2/3 and L5 Pyrs (Figure 8E-F).

Importantly, our modeling studies revealed a threshold for PV disinhibition required to generate recurrent activity between L2/3 and L5. Systematic alteration of PV input strength indicated that recurrent activity across L2/3 and L5 could be elicited when feedback inhibition from PV neurons in L2/3 was reduced by as little as 10%. Thus, although this model circuit lacks several critical elements of the intact cortical circuit, including recurrent excitation within L2/3 and also other sources of inhibition (such as from somatostatin interneurons; (Pfeffer et al., 2013; Urban-Ciecko and Barth, 2016), it successfully isolates key components that accurately reproduce experimental data. More importantly, the model indicates that reduced PV inhibition in L2/3 is necessary to permit the emergence of prolonged recurrent activity between L2/3 and L5 and that an increase of the POM input alone is not sufficient to support recurrent activity across layers.

## Discussion

Disinhibition of neural circuits has been widely proposed as a mechanism to enable excitatory synaptic plasticity. Early studies of hippocampal long-term potentiation indicated that pharmacological suppression of GABAergic transmission was required for glutamatergic synaptic strengthening (Wigström and Gustafsson, 1983). Although both anatomical and electrophysiological changes in cortical PV neuron function have been well-documented in sensory deprivation conditions (Kreczko et al., 2009; Hengen et al., 2013; Kuhlman et al., 2013; Li et al., 2014; Gainey et al., 2018), the role of PV neurons in sensory-based learning remains poorly understood. Recent studies have suggested that a *state-dependent* suppression of inhibition in cortical circuits may gate excitatory synaptic strengthening (Williams and Holtmaat, 2019) that may be important during learning (Letzkus et al., 2011; Abs et al., 2018).

Disinhibition during learning has also been reported as a persistent reduction of inhibitory synapses, albeit from an unidentified source (Donato et al., 2013; Sarro et al., 2015; Kida et al., 2016). This study identifies a precise source of GABAergic inhibition – PV neurons – that is altered in the early phases of sensory learning. Because experiments were carried out in acute brain slices and fixed tissue, the reduction in PV input described here is not simply state-dependent, but is manifested in both structural and functional changes in synaptic output. In addition, our experiments localize reduced PV inhibition specifically to superficial but not deep layers of the cortex. The laminar specificity of reduced PV inhibition may provide clues to how sensory-reward coupling may selectively engage some GABAergic circuits but not others within the cortical column.

### Anatomical and electrophysiological detection of PV input plasticity

We show that PV-mediated inhibition of L2/3 Pyr neurons is highly sensitive to reward-based sensory-association training, decreasing rapidly at the onset of training and returning to baseline levels as behavioral performance plateaus. Electrophysiological and anatomical reduction in PV inputs showed a striking correlation. The 35% reduction in mean PV-ChR2 IPSC amplitude in L2/3 Pyr neurons after 24 hours of SAT, closely matched the ~39% reduction in PV-associated synapses at the soma/dendrites. Anatomical analyses of PV inputs during SAT revealed that PV synapse loss could be observed at the soma as well as at higher-order dendrites, further underscoring recent quantitative data indicating that PV inputs are broadly distributed across Pyr neurons (Kubota et al., 2015; Kuljis et al., 2019).

PV neurite alignment with FAPpost-labeled postsynaptic sites indicated that PV synapse loss was associated with the removal of postsynaptic structures. Interestingly, a comparison of SAT-dependent changes in PV neurite apposition with L2/3 Pyr neurons, without the requirements of a postsynaptic marker present, did not reveal reductions in presynaptic PV+ structures after 24 hours of SAT. Combined with the finding that PV input returned to control levels after prolonged training, these anatomical data suggest that the inhibitory synapse plasticity described here is accompanied by the dismantling of post-synaptic structures, not by the large-scale movement or elimination of PV axons and/or release sites.

### Temporal control of inhibition during learning

L2/3 Pyr neurons undergo a reduction in PV input that is initiated rapidly after the onset of SAT. The selective decrease in PV input observed during reward-associated sensory training but not passive sensory exposure suggests that reward signals are integrated in S1 and facilitate the removal of PV synapses. How might this be implemented?

PV neurons are responsive to cholinergic signaling (Kruglikov and Rudy, 2008; Letzkus et al., 2011) and are embedded in a complex and highly organized network of molecularly defined inhibitory neurons in the neocortex (Pfeffer et al., 2013; Jiang et al., 2015; Barth et al., 2016). Cell-type specific recordings in sensory cortex indicate that reinforcement cues can acutely suppress PV neuron activity (Letzkus et al., 2011), possibly related to cholinergic activation of L1 and/or vasoactive intestinal peptide (VIP)-expressing interneurons (Arroyo et al., 2012; Chen et al., 2015a). Short-term suppression of PV activity, experienced over multiple stimulus-reward pairings during SAT, may trigger the structural and functional changes to PV synapses observed in this study. It remains unknown how interactions between other types of inhibitory neurons are changed during sensory learning, and it is likely that other types of inhibitory neurons are at least acutely engaged during sensory-evoked plasticity (Abs et al., 2018; Yaeger et al., 2019). It will be critical for future studies to determine whether other neuron types show SAT-associated reductions in PV inhibition, or whether PV-disinhibition is restricted to a subset of L2/3 Pyr neurons, for example those defined by projection target (Chen et al., 2016).

What are the consequences of decreased PV input to L2/3 Pyr neurons? Reduced PV inhibition, in combination with the strengthening of excitatory thalamic and intracortical synaptic pathways (Audette et al., 2019), may increase stimulus-evoked and also prolonged cortical activity in the early stages of SAT. Under baseline conditions, L2/3 Pyr neurons are only weakly driven by sensory input at short latencies (O’Connor et al., 2010; Lefort and Petersen, 2017), and they receive potent feedback PV inhibition that can be driven by the firing of a single Pyr neuron (Jouhanneau et al., 2018). The sparse firing of Pyr neurons in superficial layers, particularly in somatosensory cortex, is likely related to this pronounced inhibition (Barth and Poulet, 2012). Our modeling studies indicate that SAT-dependent reductions in PV input to L2/3 Pyr neurons are sufficient to enable enhanced recurrent activity within and between layers that is associated with reinforcement learning (Audette et al., 2019). These data provide insight into the powerful role that inhibition in L2/3 can play in influencing cortical output during both sensory-evoked and spontaneous activity (Vogels et al., 2011; Wilmes and Clopath, 2019). Stimulus-initiated recurrent activity in PV-disinhibited superficial layers of cortex may be important in the formation of new adaptive connections that link whisker stimulation with reward during association learning.

### Layer-specific PV plasticity

Our earlier study showed that after 24 hrs of SAT, thalamocortical (POm) input potentiation has occurred L5 Pyr neurons. If disinhibition was critical for this excitatory synaptic plasticity, we expected it would be manifested in L5 Pyr neurons, since PV-mediated inhibition is a prominent feature of sensory processing circuitry in L5 (Jiang et al., 2015). PV neurons in L5 are a potent source of thalamic feedforward inhibition onto neighboring L5 Pyr neurons (Audette et al., 2017), and L5 Pyr neurons show larger PV-IPSCs than their L2/3 counterparts. However, the lack of PV input change at 12 and 24 hours of SAT suggests that L5 Pyr disinhibition was not detected because it had already renormalized. Instead, our findings suggest thalamocortical plasticity in L5 Pyr neurons may not require a reduction in PV input, possibly because at baseline they typically show higher firing rates that may be sufficient for plasticity induction during learning (De Kock et al., 2007; Audette et al., 2017). Alternatively, it is possible that disinhibition of L5 Pyr neurons is initiated by SAT, but is state-dependent (Kruglikov and Rudy, 2008) or is implemented through a different inhibitory source such as SST cells, particularly at the apical tuft of the L5 Pyr dendrite.

Where within the cortical column are the PV neurons that are altered during SAT? Our anatomical and electrophysiology analyses could not reveal the laminar location of PV neurons that reduced their output to L2/3 neurons. Experimental data indicate that PV neurons across all layers innervate L2/3 Pyr neurons, whereas L5 Pyr neurons receive most of their inhibition from infragranular neurons (Kätzel et al., 2011; Pfeffer et al., 2013; Jiang et al., 2015; Kubota et al., 2015; Barth et al., 2016; Frandolig et al., 2019). Future experiments characterizing SAT-related PV disinhibition across layers will help determine whether plasticity is restricted to PV neurons that reside in a particular layer, and will ultimately help determine the circuit and synaptic requirements that initiate PV-mediated disinhibition during learning.

## Conclusion

We hypothesize that PV disinhibition may be a necessary step in driving brain plasticity associated with long-lasting behavioral change during learning (Barth and Ray, 2019). Importantly, our study indicates that PV neurons can differentiate between incidental and meaningful sensory information as their plasticity was selectively engaged only during reward association training, suggesting that they are a critical node in learning-associated changes in the cortical circuit. Cortical neurons in primary sensory cortex are well-positioned to receive and amplify delayed, reward-related cues that facilitate excitatory synaptic remodeling and activation of downstream brain areas that are directly linked to behavioral change. Thus, the disinhibition of L2/3 Pyr neurons may be a key step in altering the response properties of cortical neurons during learning.

## Methods

### Animals

All experimental procedures were conducted in accordance with the NIH guidelines and approved by the Institutional Animal Care and Use Committee at Carnegie Mellon University. For functional assessment of PV-to-Pyr synaptic strength, Cre-dependent channelrhodopsin-2 (ChR2; Ai32 strain Jackson Lab Stock ID 012569; Madisen et al., 2012) and PV-Cre (Jackson Lab Stock ID 008069; Hippenmeyer et al., 2005) double-transgenic knock-in mice were used (male and female, postnatal day (P)25-29). For a subset of experiments examining PV excitability, PV neurons were targeted using PV-tdTom mouse line (Jackson Lab Stock ID 027395).

For anatomical experiments, PV-Cre and Cre-dependent YFP (Ai3 strain Jackson Lab Stock ID 007903; Madisen et al., 2010) double-transgenic knock-in mouse (male and female) barrel cortex was stereotaxically injected with FAPpost (0.1uL), a neuroligin1-based rAAV construct that mediates far-red fluorescent signal at postsynaptic sites (Kuljis et al., 2019). Virus was introduced through a small craniotomy (from bregma: x=−3, y=−0.9, z=−0.5 mm) using a Nanoject II (Drummond Scientific Company; Broomall, PA) in isoflurane anesthetized mice at P15-17. Six to 8 days later, animals underwent whisker-stimulation reward association training.

### Automated sensory association training

We used an automated, high-throughput experimental paradigm for gentle airpuff-reward training for sensory learning as described previously (Audette et al., 2019). Briefly, animals were housed in modified homecages equipped with an SAT chamber in which initiating nosepokes at the waterport caused an infrared beam break that triggered trial onset with a random variable delay (0.2-0.8s) preceding the conditioned stimulus. During SAT, 80% of (stimulus) trials began with administration of a gentle, downward-projecting airpuff directed against right-side whiskers (4-6 PSI, 0.5 s duration). One second after trial onset, a water reward (~8-25 µL) was delivered to the lickport. For the remaining 20% of (blank) trials, nosepokes triggered an approximately 2-3 second timeout (depending on random delay duration; Figure 1). During pseudotraining, airpuff stimulation was administered in 80% of (stimulus) trials, and water was delivered for half of those trials. For the remaining 20% of (blank) trials, water was delivered for half of those trials. Thus, in SAT experiments, airpuff was predictive of water reward, and in pseudotraining experiments, sensory stimulation was uncoupled from water-reward. For SAT and pseudotraining experiments, litter and cage-matched controls were used. Performance was calculated as the difference in anticipatory lick rates (0.7-1 ms following trial onset) for stimulus trials vs. blank trials (Lick_Water_-Lick_Blank_). Mean anticipatory lick rates for each animal were calculated in 4-hour bins, from SAT-chamber acclimation 24 hours before experiment onset through to the end of the experiment. For reliable estimates of performance, we required that a minimum of 10 total trials (stimulus and blank trials) within a 4-hour window had to be completed for an animal’s data to be included.

### Electrophysiology

At midday (11am-2pm) following SAT (SAT24 or SAT120) or housing in training cages without airpuff exposure (Ctrl24 or Ctrl120), mice (P25-29) were briefly anesthetized with isoflurane before decapitation. Angled-coronal slices (45° rostro-lateral; 350 µm thick) designed to preserve columnar connections in somatosensory cortex were prepared in ice-cold artificial cerebrospinal fluid (ACSF) composed of (in mM): 119 NaCl, 2.5 KCl, 1 NaH_2_PO_4_, 26.2 NaHCO_3_, 11 glucose, 1.3 MgSO_4_, and 2.5 CaCl_2_ equilibrated with 95%CO_2_/5%O_2_. Slices were allowed to recover at room temperature in ACSF for one hour in the dark before targeted whole-cell patch-clamp recordings were performed using an Olympus light microscope (BX51WI) and borosilicate glass electrodes (4-8 MΩ resistance) filled with internal solution composed of (in mM): 125 potassium gluconate, 10 HEPES, 2 KCl, 0.5 EGTA, 4 Mg-ATP, 0.3 Na-GTP, and trace amounts of AlexaFluor 594 (pH 7.25-7.30, 290 mOsm). Because of the need to verify cell type identity using action potential waveform, we used a K-gluconate based internal solution.

Electrophysiological data was acquired using a MultiClamp 700B amplifier, digitized with a National Instruments acquisition interface, and collected using MultiClamp and IgorPro6.0 software with 3kHz filtering and 10 kHz digitization. L2/3 and L5 Pyr neurons were targeted based on Pyr morphology, using the pial surface and dense PV-Ai32 fluorescence in L4 barrels for laminar orientation.

Following whole-cell break in, presumptive Pyr cell identity was confirmed based on hyperpolarized resting membrane potential (approximately −70mV in L2/3 and −60mV in L5), input resistance (approximately 100-200 MΩ; < 400MΩ cut-off), and regular-spiking (RS) action potential waveforms recorded in responses to progressive depolarizing current injection steps recorded in current-clamp mode (50-400 pA, Δ50 pA steps, 0.5s duration). L5 Pyr neurons were typically in the top to middle portion of L5 (L5a) and had either a RS or intrinsically bursting phenotype with current injection. Only cells with a stable baseline holding potential, resting membrane potential <-50mV, and access resistance <40MΩ were analyzed. PV-mediated inhibitory postsynaptic currents (IPSCs) were isolated as previously described (Pfeffer et al., 2013; Vickers et al., 2018). Blue light stimulation was used to evoke PV-IPSCs (470nm, 0.48mW LED, 5 ms pulse). Consistent with a chloride-mediated current, the reversal potential for optically-evoked currents was experimentally determined to be −78 ± 4 mV. Five minutes after break-in, Pyr cells were voltage-clamped (VC) at −50 mV and PV-mediated IPSCs were collected, where peak amplitude was calculated from the average of 10 sweeps (0.1 Hz). For recordings where a single light-pulse evoked multiple IPSC peaks, only the amplitude of the first peak was measured. For a subset of cells, picrotoxin (50µM) was applied to confirm optically-evoked IPSCs were GABA_A_ receptor-mediated, and in all cases picrotoxin abolished hyperpolarizing outward currents. For a subset of experiments, recordings were performed blinded to the experimental condition. In some cases, optically-evoked currents were measured in parallel by a separate experimenter using a different electrophysiology rig and blue-light optical filter. Illumination intensity was calibrated between rigs using average PV-IPSC response in control animals. Across experiments, responses collected on each rig were not significantly different so all data were pooled in the final analysis.

To assess L2/3 PV neuron excitability, PV neurons were targeted in either PV-Cre × Ai32 or PV-tdTom transgenic mouse tissue for current-clamp recordings (Barth et al., 2004). PV neuron identity was verified by reporter fluorescence, fast-spiking phenotype in response to direct depolarizing current injection, and/or the presence of excitatory photocurrents in response to blue light stimulation. Only PV cells with a stable baseline holding potential and resting membrane potential < −45mV were analyzed.

### Anatomy

At mid-day following 24 hours of SAT, animals were anesthetized with isoflurane and transcardially perfused using 20mL PBS (pH 7.4) followed by 20mL 4% paraformaldehyde (PFA) in PBS (PFA; pH 7.4). Brains were removed, and postfixed overnight at 4°C in 4% PFA before transfer into 30% sucrose cryoprotectant. After osmotic equilibration, 45 µm-thick brain sections were collected using a freezing-microtome. Free-floating brain sections containing dTom-expressing cells in the barrel cortex were washed with PBS before 30-minute room temperature incubation with MG-Tcarb dye (300nM in PBS) for activation of the far-red fluorescence of the FAP (Pratt et al., 2017).

Pyr neurons were identified by their pyramid-shaped cell body, a narrow axon descending from the base of their soma, a prominent apical dendrite and laterally projecting, spiny basal dendrites. Confocal image stacks centered around a well-isolated, FAPpost-expressing Pyr soma were collected with a LSM 880 AxioObserver Microscope (Zeiss) using a 63x oil-immersion objective lens (Plan-Apochromat, 1/40 Oil DIC M27) with the zoom factor set to 1 and the pinhole set at 1.0 Airy disk unit for each fluorescence channel. Optimal laser intensities for each channel were set to avoid pixel saturation for each cell independently. Fluorescence acquisition settings were as follows: YFP (excitation λ514, detection λ517–535), dTom (excitation λ561, detection λ561–597), and MG/FAP (excitation λ633, detection λ641– 695). Maximum image size was 1024×1024 pixels, to collect 135 × 135 × ≤ 45µm images, with corresponding 0.13 × 0.13 × 0.3µm voxel dimensions.

Synapse distribution analysis was carried out using previously published methods for the FAPpost synaptic marker (Kuljis et al., 2019). In brief, after Carl Zeiss image files were imported into Imaris (v8.4 with FilamentTracer; Bitplane; Zürich, Switzerland), the dTom cell fill was used to create a 3D Pyr neuron rendering using Imaris macros to create a combination of “surface” and “filament” objects. FAPpost puncta were then reconstructed as “surfaces” using an estimated 0.5µm diameter, 4-voxel minimum, and spit-touching object setting using the same 0.5µm diameter. FAPpost “surfaces” were digitally assigned to a given neuron if their edges lay within 0.5µm of the soma surface (inner and outer edge), or ≤1µm from dendrite. Puncta “surfaces” were converted into puncta “spots” (created using automatic intensity-maxima background-subtraction thresholds with an estimated 0.5µm diameter) using “surface” object centroids. Presynaptic neurite reconstructions were created using automatic background-subtraction thresholding of presynaptic PV-YFP fluorescence using an estimated diameter of 0.6µm, split-touching object diameter threshold of 1µm (applied with automatic “quality” filter setting), and a 1µm^2^ minimum surface area. To digitally correct for z-axis related signal drop-off, neurite reconstruction using automatic settings were generated separately for every 10µm of z-depth resulting in similar density and size profiles for both superficial and deep presynaptic neurite reconstructions. Finally, FAPpost puncta “spots” were assigned as PV+ using a distance threshold of 0.15µm from spot centroid to presynaptic neurite edge (PV synapse).

Since discrete classes of PV neurons may differentially target Pyr neuron compartments (Kubota et al., 2016; Vereczki et al., 2016; Feldmeyer et al., 2017; Lu et al., 2017), compartment-specific methods for assessing PV synapses were used to serve as a guide for evaluating whether a specific population of presynaptic PV neurons might be differentially affected by SAT. During preliminary analysis, PV synapse density across Pyr dendrites was assessed separately for apical and basal dendrite segments (across branch orders), soma, and axon compartments by taking the total number of PV-assigned synapses for each compartment and dividing it by the total length (for dendrites and axon) or surface area (for soma). Since a similar decrease in PV-assigned synapse density was observed across all dendritic compartments (low and higher order apical and basal dendrites), all dendritic compartments were pooled in the final analysis, and reported densities were calculated using the total number of spots (total FAPpost and PV synapses) divided by total length of dendrite analyzed.

### Computational modeling

We generated a simplified network consisting of 5 neurons to capture the minimal elements of the cortical circuit engaged by POm activation. This included a POm neuron connected to L2 Pyr, L5 Pyr, and L5 PV neurons; a L2 Pyr and a L2 PV neuron with reciprocal connectivity; and a L5 Pyr and a L5 PV neuron with reciprocal connectivity. We stimulate the POm neuron at time t = 6 [a.u] with an amplitude of POm=1 [a.u]. Pyr neurons are modelled as Integrate-and-Fire (Gerstner and Kistler, 2002), such that their voltages, *v_L5_* for L5 and *v_L2_* for L2, can be written as:

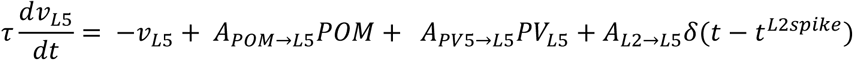

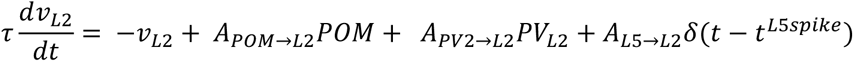

where τ = 10[*a.u*] is the membrane time constant, the synaptic strength are denoted by the different As,*δ* is the Dirac delta function and *t^L2spike^* and *t^L5spike^* are the times of the L2 and L5 Pyrs respectively with a 2 [a.u] time delay. The Integrate-and-Fire neuron is spiking when the voltage crosses the threshold of 0.015 [a.u.] and the voltage is then set back to 0. L2 and L5 Pyrs are mutually exciting with an amplitude of *A_L2→L5_* = *A_L5→L2_* = 0.16 [a.u.], as suggested by experimental data (Jiang et al., 2015; Barth et al., 2016; Lefort and Petersen, 2017). In accordance with experimental measurements (Audette et al., 2019), POm input drives L5 Pyr neurons 5 times more strongly than L2 neurons so that *A_POM→L5_* = 0.5[a.u] and *A_POM→L2_* = 0.1 [a.u]. The PV neurons are simply modelled as linear integrator of currents that summate the pyramidal cell current. Based on experimental data (Audette et al., 2017), *PV_L5_* but not *PV_L2_* receive POM input with an amplitude of 0.05 [a.u.]. PV neurons inhibit within-layer pyramidal cells with an amplitude of *A_PV5→L5_* = *A_PV2→L2_* = 5 (Avermann et al., 2012; Jiang et al., 2015; Barth et al., 2016; Litwin-Kumar et al., 2016). Network simulations proceed for 30 [a.u] amount of time. To simulate conditions after 24 hrs of SAT, we increased the POm input to L5 pyramidal cell, *A_POM→L5_*, by 20% (Audette et al., 2019), we decrease the *PV_L2_* amplitude to L2 pyramidal cell, *A_PV2→L2_* by 40% and we perform, as before, one single stimulation of amplitude of POm=1 [a.u.] at time t = 6 [a.u.].

### Statistics

Mean anticipatory lick-rate and performance (±SEM) for each 4-hour time bin was used to represent average group behavior. PV-IPSC magnitudes, membrane potential, input resistance, rheobase current, optically evoked spike count, as well as PV neurite, PV synapse, and FAPpost densities for dendrite (per µm) and soma (per µm^2^) across Pyr or PV neurons was assessed for statistical significance using the Mann-Whitney U test (GraphPad Prism, v7; San Diego, CA). Comparisons were made between 24 hr control and SAT groups, 120 hr control and SAT groups, and 24 hrs pseudotraining control and pseudotraining groups within layer (L2/3 or L5). IPSC amplitudes are reported in text and represented in graphs as mean±SEM. Unless otherwise noted, excitability measures, PV neurite, and PV synapse densities averaged by cell are reported in text and represented in graphs as mean±SD (with individual cell values overlaid). Effect of current injection step and experimental condition on firing frequency responses was assessed using two-way ANOVA (OriginPro, Northampton, MA). Statistical significance, *p*<0.05.

## Acknowledgements

Special thanks to Joanne Steinmiller for expert management of transgenic mice, Sarah Bernhard for SAT cage design and technical support, Ajit Ray and Alex Hsu for custom MatLab scripts for behavioral analysis, Marcel Bruchez for providing reagents for FAPpost detection, and members of the Barth Lab for helpful comments on the manuscript.

---

The following figure supplements are available for figure 2: **Source Data 1.** SAT24 PV IPSC statistics table.

**Figure 2-source data 1.** SAT24 PV IPSC Statistics Table

| *Layer* | *Condition* | *n*<br>*cells* | *Median PV*<br>*IPSC (pA)* | *Mann*<br>*Whitney U* | *p* |
| --- | --- | --- | --- | --- | --- |
| Layer 2/3 | Ctrl24 | 19 | 363 | 69.5 | 0.0028 |
|  | SAT24 | 17 | 227 |  |  |
| Layer 5 | Ctrl24 | 13 | 731 | 89 | 0.38 |
|  | SAT24 | 17 | 672 |  |  |

The following figure supplements are available for figure 3: **Source Data 1.** SAT120 PV IPSC statistics table.

**Figure 3-source data 1.** SAT120 PV IPSC Statistics Table

| *Layer* | *Condition* | *n*<br>*cells* | *Median PV*<br>*IPSC (pA)* | *Mann*<br>*Whitney U* | *p* |
| --- | --- | --- | --- | --- | --- |
| Layer 2/3 | Ctrl120 | 25 | 310 | 452 | 0.52 |
|  | SAT120 | 40 | 365 |  |  |
| Layer 5 | Ctrl120 | 27 | 646 | 12 | 1.0 |
|  | SAT120 | 19 | 578 |  |  |

The following figure supplements are available for figure 4: **Source Data 1.** Pseudo24 PV IPSC statistics table.

**Figure 4-source data 1.**
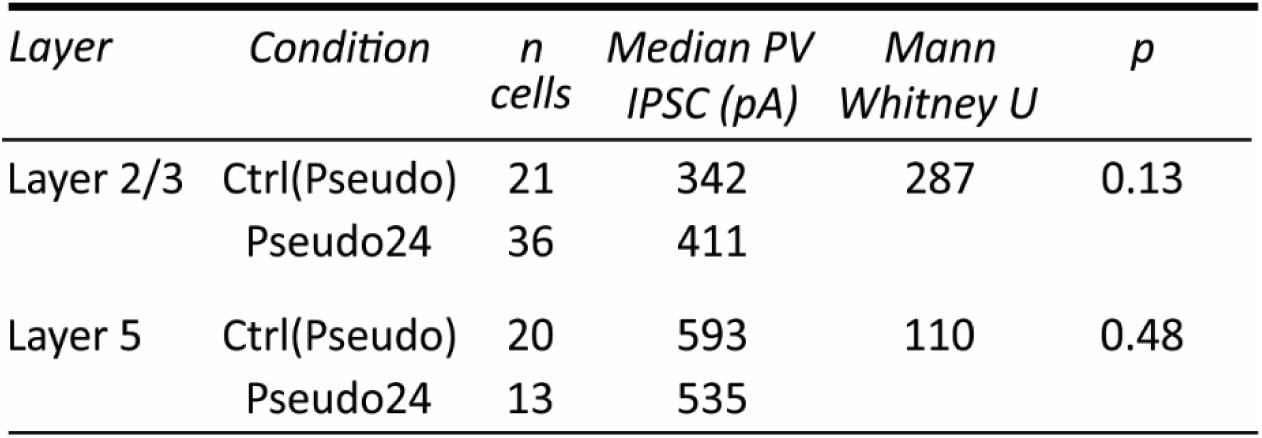
Pseudo24 PV IPSC Statistics Table

| *Layer* | *Condition* | *n*<br>*cells* | *Median PV*<br>*IPSC (pA)* | *Mann*<br>*Whitney U* | *p* |
| --- | --- | --- | --- | --- | --- |
| Layer 2/3 | Ctrl(Pseudo) | 21 | 342 | 287 | 0.13 |
|  | Pseudo24 | 36 | 411 |  |  |
| Layer 5 | Ctrl(Pseudo) | 20 | 593 | 110 | 0.48 |
|  | Pseudo24 | 13 | 535 |  |  |

The following figure supplements are available for figure 5: **Source Data 1.** L2/3 PV neuron excitability summary statistics table.

**Figure 5-source data 1.** L2/3 PV Neuron Excitability Summary Statistics

**Figure 5—source 1. L2/3 PV Neuron Excitability Summary Statistics**
| *Measure*<br>*(units)* | *Condition* | *n*<br>*cells* | *Mean±SD* | *Median* | *Mann*<br>*Whitney U* | *p* |
| --- | --- | --- | --- | --- | --- | --- |
| Vm<br>(mV) | Ctrl24 | 13 | -63±8 | -63 | 61.5 | 0.58 |
|  | SAT24 | 11 | -62±6 | -62 |  |  |
| Rin<br>(MΩ) | Ctrl24 | 12 | 410±235 | 308 | 58 | 0.64 |
|  | SAT24 | 11 | 408±271 | 290 |  |  |
| Optically<br>evoked<br>spikes (#) | Ctrl24 | 12 | 1.3±1.2 | 1 | 32.5 | 0.75 |
|  | SAT24 | 6 | 1.3±0.8 | 1 |  |  |
| Rheobase<br>(pA) | Ctrl24 | 12 | 71±36 | 69 | 49 | 0.49 |
|  | SAT24 | 10 | 89±55 | 78 |  |  |

The following figure supplements are available for figure 6: **Source Data 1**. SAT24 PV synapse density statistics table. **Figure Supplement 1**. Fluorescence-based analysis approach for input-specific synapse mapping using Imaris.

**Figure 6-source 1.**
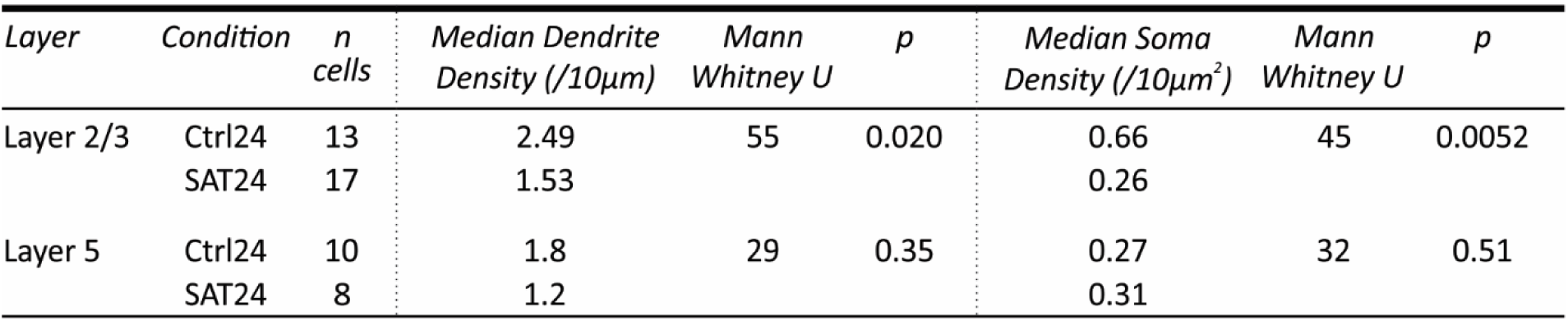
SAT24 PV Synapse Density Statistics Table

| *Layer* | *Condition* | *n*<br>*cells* | *Median Dendrite*<br>*Density (/10µm)* | *Mann*<br>*Whitney U* | *p* | *Median Soma*<br>*Density (/10µm<sup>2</sup>)* | *Mann*<br>*Whitney U* | *p* |
| --- | --- | --- | --- | --- | --- | --- | --- | --- |
| Layer 2/3 | Ctrl24 | 13 | 2.49 | 55 | 0.020 | 0.66 | 45 | 0.0052 |
|  | SAT24 | 17 | 1.53 |  |  | 0.26 |  |  |
| Layer 5 | Ctrl24 | 10 | 1.8 | 29 | 0.35 | 0.27 | 32 | 0.51 |
|  | SAT24 | 8 | 1.2 |  |  | 0.31 |  |  |

**Figure 6–supplement 1.**
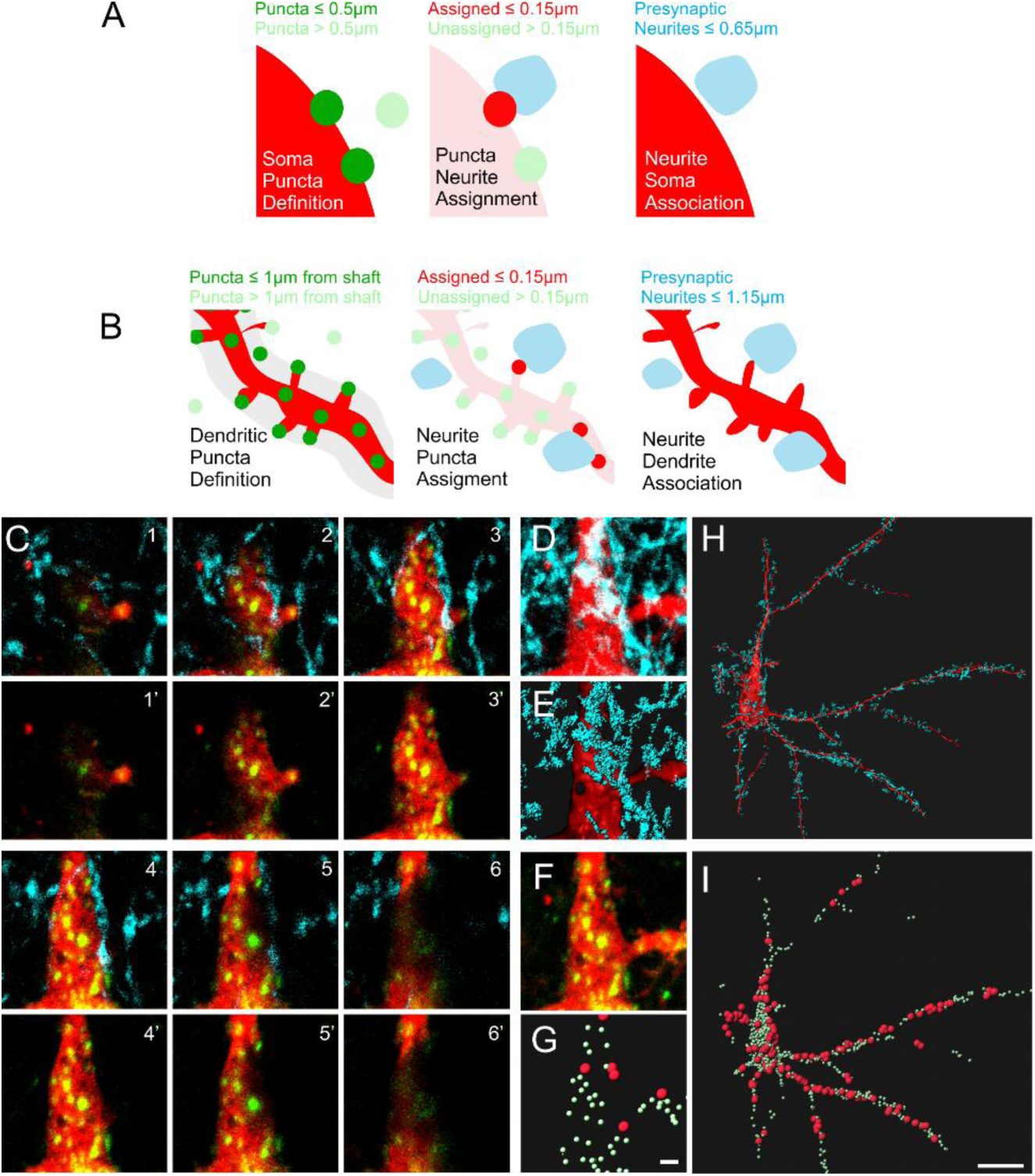
Fluorescence-based analysis approach for input-specific synapse mapping using Imaris. (**A**) Distance threshold parameters for input assignment on FAPpost puncta on soma (**B**) and dendrite. (**C**) Six serial optical sections in a L2/3 Pyr neuron primary apical dendrite labeled with FAPpost (synaptic sites in green, dTom cytoplasmic fill in red; panels 1’-6’). Panels 1-6 show overlay with presynaptic PV-YFP neurites (cyan). (**D**) Flattened stack of the region in *C*. (**E**) 3D rendering of PV neurites (cyan) and Pyr dendrite (red) in *D*. (**F**) As in D, but for FAPpost and dTom. (**G**) PV-assigned FAPpost (large red) and unassigned FAPpost synapses (small green). Scale bar=1µm. (**H**) As in *E*, but for a larger region showing PV neurite contacts on soma and a subset of the dendritic arbor. (**I**) As in *H*, but for PV-assigned (red) and unassigned (green) FAPpost synapses. Scale bar =20µm.

The following figure supplements are available for figure 7: **Source Data 1.** SAT24 PV neurite crossing density statistics table.

**Figure 7-source 1.** SAT24 PV Neurite Crossing Density Statistics Table

| *Layer* | *Condition* | *n*<br>*cells* | *Median Dendrite*<br>*Density (/10µm)* | *Mann*<br>*Whitney U* | *p* | *Median Soma*<br>*Density (/10µm<sup>2</sup>)* | *Mann*<br>*Whitney U* | *p* |
| --- | --- | --- | --- | --- | --- | --- | --- | --- |
| Layer 2/3 | Ctrl24 | 13 | 1.4 | 89 | 0.39 | 1.8 | 91 | 0.43 |
|  | SAT24 | 17 | 1.2 |  |  | 1.5 |  |  |
| Layer 5 | Ctrl24 | 10 | 2 | 31 | 0.46 | 2.1 | 37 | 0.83 |
|  | SAT24 | 8 | 1.9 |  |  | 2.1 |  |  |

